# Renal tubular function and morphology revealed in kidney without labeling using three-dimensional dynamic optical coherence tomography

**DOI:** 10.1101/2023.05.01.539010

**Authors:** Pradipta Mukherjee, Shinichi Fukuda, Donny Lukmanto, Thi Hang Tran, Kosuke Okada, Shuichi Makita, Ibrahim Abd El-Sadek, Yiheng Lim, Yoshiaki Yasuno

## Abstract

Renal tubule has distinct metabolic features and functional activity that may be altered during kidney disease. In this paper, we present label-free functional activity imaging of renal tubule in normal and obstructed mouse kidney models using three-dimensional (3D) dynamic optical coherence tomography (OCT) *ex vivo*. To create an obstructed kidney model, we ligated the ureter of the left kidney for either 7 or 14 days. Two different dynamic OCT (DOCT) methods were implemented to access the slow and fast activity of the renal tubules: a logarithmic intensity variance (LIV) method and a complex-correlation-based method. Three-dimensional DOCT data were acquired with a 1.3 μm swept-source OCT system and repeating raster scan protocols. In the normal kidney, the renal tubule appeared as a convoluted pipe-like structure in the DOCT projection image. Such pipe-like structures were not observed in the kidneys subjected to obstruction of the ureter for several days. Instead of any anatomical structures, a superficial high dynamics appearance was observed in the perirenal cortex region of the obstructed kidneys. These findings suggest that volumetric DOCT can be used as a tool to investigate kidney function during kidney diseases.

## Introduction

The kidney is a complex organ that performs several vital tasks including filtering the blood, removing toxins, and maintaining an acid-base balance. The functional unit of the kidney is the nephron, which is composed of renal corpuscles (glomerulus with Bowman’s capsule) and tubular segments^1^. The renal tubular compartment consists of a proximal convoluted tubule (PCT), an intermediate tubule (loop of Henle), and a distal convoluted tubule (DCT). The renal tubules (mainly PCT and DCT) are responsible for metabolic needs such as fatty acid oxidation, glucose metabolism, and amino acid metabolism^2–6^.

The kidney can be acutely injured by ureter obstruction, ischemia, toxins, drugs, and infection, or can be chronically damaged by diabetes mellitus or hypertension^7,8^. The renal tubules are vulnerable to both acute and chronic kidney injuries, and it is known that during such injuries, renal tubular metabolism is altered or even damaged^9^. To understand the metabolism of healthy renal tubules and how it changes during kidney injury, it is necessary to investigate the functional activity of renal tubules using small animal models. As the renal tubule extends from the renal cortex to the deep region and has a three-dimensional architecture, it is important to visualize the functional activity of deep renal tissue without any slicing.

Two-photon microscopy (TPM) allows researchers to visualize and understand dynamic intracellular and intercellular activities in kidney tissue^10–12^. Hence, it is the preferred experimental tool for investigating kidney physiology and diseases. TPM uses fluorescent probes to visualize structural and functional changes in the kidney. Due to the use of fluorescent probes, the acquired data of TPM are compromised by photobleaching and phototoxicity. Additionally, TPM has a limited imaging penetration depth of around 150 μm^12^. This limited penetration depth poses a challenge for visualizing the functional activity of deep renal tubule compartments. Therefore, an imaging technique that can accurately assess both the structural and functional activity of renal tubules without the use of exogenous contrast agents is required.

Optical coherence tomography (OCT) is a label-free, cross-sectional, and three-dimensional imaging modality that uses low-coherence interferometry principles to reconstruct tomographic images of tissue structures^13^. OCT has resolution down to a few micrometers and imaging penetration of a few millimeters (around 1 – 2 mm). Several studies demonstrated the ability of OCT imaging to visualize three-dimensional internal microstructures of the kidney, such as renal tubules and glomerulus either *ex vivo* or *in vivo*^14,15^. Although OCT is particularly suitable for examining renal tissue, it can only visualize its static property, such as renal morphology. The functional activity of renal tubule compartments cannot be accessed using conventional OCT.

This limitation of conventional OCT, i.e., functional activity investigation can be overcome using extended OCT contrast so-called dynamic OCT (DOCT). Using DOCT, it is possible to visualize functional tissue structures in living organisms at a cellular or sub-cellular scale without the need for contrast agents^16–22^. This is achieved by acquiring several OCT frames with millisecond temporal resolution over a period of several seconds, followed by statistical analysis of the time-sequence signal. It was first introduced using a time-domain full-field OCT which generates *en face* dynamics image^16^ and was later implemented in Fourier-domain OCT that generates cross-sectional (B-scans) dynamics image^18,19^. Three-dimensional visualization of dynamics imaging has also become possible using ultra-fast OCT systems^17,20^ or long measurement time^23,24^. Recently, we demonstrated a custom-designed scanning protocol that uses a standard-speed swept-source OCT system to acquire volumetric tomography of tissue dynamics within a few seconds^25^.

Several DOCT methods for visualizing the functional dynamic structures of tissue have recently been demonstrated. These methods can be categorized into magnitude^18,21,26^, frequency^16,17,19^, and time-correlation analysis^18,27,28^ of OCT signal fluctuation. Among the magnitude methods, logarithmic intensity variance (LIV) contrasts the fluctuation magnitude of the OCT signal intensity^18^. In our previous studies, LIV successfully revealed high-contrast dynamic structures in *ex vivo* mouse organs^29,30^ and *in vitro* tumor spheroids^18,25^. Among the time-correlation methods, Lee *et al*. used complex correlation analysis of time-sequential OCT signals to extract the velocity and diffusion coefficient of moving particles^27^. In general, the complex correlation-based algorithm is suitable for visualizing fast (millisecond-scale) tissue activity, whereas LIV is sensitive to relatively slow (second-scale) functional activity. Hence, a combination of LIV and complex correlation-based methods may provide a more comprehensive understanding of renal tubule dynamics.

In this paper, we demonstrate three-dimensional renal tubular functional activity imaging using DOCT. Two different signal processing algorithms and two custom-designed scanning protocols were used to access the slow and fast activity of renal tubule. For the slow functional activity imaging, the LIV of time-sequence OCT frames was calculated^18,25^ within a 6.12-s time window with an inter-frame time interval of 409.6 ms. For the fast activity visualization, complex correlation of time sequence OCT frames^31^ was computed within a 39-ms time window with an interframe time interval of 12.8 ms. Here, we refer to the correlation-based dynamic OCT contrast as ‘fast dynamic OCT (Fast-DOCT)’. DOCT imaging was performed using our custom-made standard-speed (50,000 A-lines/s) swept-source OCT system^32,33^. Two different kidney model tissues were studied, involving normal and obstructed kidney models. For the obstructed kidney model, the ureter of the left kidney was obstructed for either 7 or 14 days. DOCT visualizes differences in the functional activity of morphological renal tubules between normal and obstructed kidney models.

## Results

We used swept-source OCT microscopy to capture the dynamics in renal tissues. The OCT microscope used was reported previously (Methods section)^32,33^. To capture the dynamic speckle patterns in live renal tissues, we recorded 16 frames at a single location within the sample, with an interframe time interval of 409.6 ms over a time window of 6.14 s. The LIV image is reconstructed by calculating the magnitude of the speckle fluctuation, which is determined by analyzing the temporal variance of the time-sequence logarithmic (dB) scale OCT intensity signal within a 6.14-s time window (see the Methods section and Section 2.2.1 of Ref.^18^ for detailed derivation). Fast-DOCT was obtained by complex correlation analysis^31^ among four repeated frames acquired with an inter-frame time separation of 13 ms within a time window of 39 ms.

### Label-free dynamics imaging of fresh normal kidney

Figure 1 displays three-dimensional OCT intensity, LIV, and Fast-DOCT images of a normal mouse kidney for a 3 mm × 3 mm lateral scanning range. The depth location of the *en face* slices and the location of the B-scans in the *en face* images are shown by the horizontal lines in the OCT intensity images [Figs. 1(a) and (b)].

**Figure 1.**
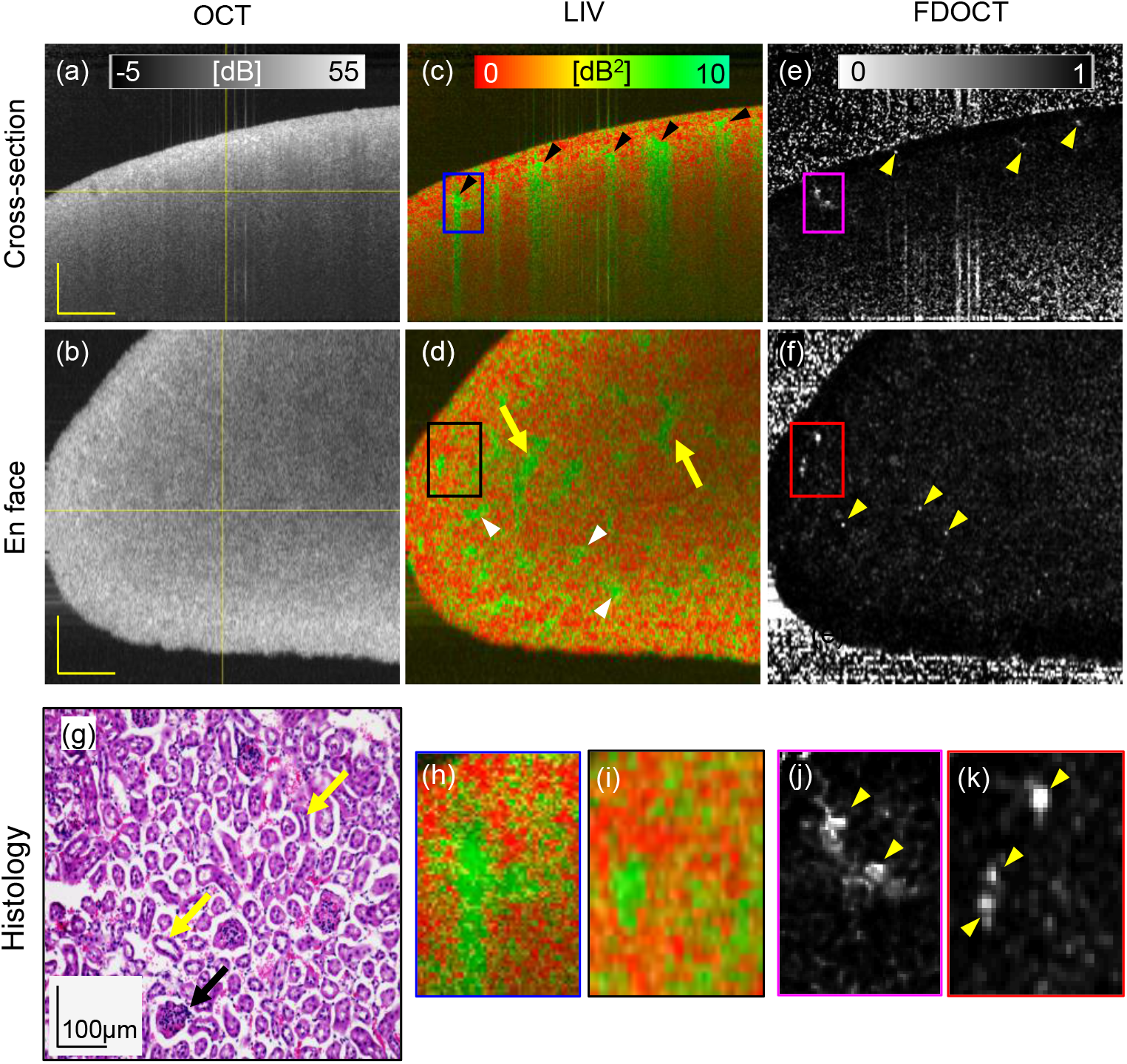
Dynamic OCT imaging of a fresh normal mouse kidney for a 6 mm × 6 mm field of view. Cross-sections from (a) scattering OCT, (c) LIV, and (e) Fast-DOCT imaging; (b, d, f) *En face* slices of the OCT, LIV, and Fast-DOCT images at the depth location indicated by the horizontal line in (a); (g) H&E stained histological micrograph; (h, j) Magnified images of the LIV and Fast-DOCT cross-section at the region indicated by the rectangular box in (c, e); (i, k) Magnified images of the LIV and Fast-DOCT *en face* slice at the region indicated by the rectangular box in (d, f). The black arrow in (g) indicates the glomerulus and the yellow arrows indicate the renal tubules of the kidney tissue. LIV: log-intensity variance; Fast-DOCT: fast dynamic OCT; H&E: Hematoxylin and eosin. Scale bar: 500 μm.

In the standard OCT intensity, the structure of the renal tissue cannot be identified and the tissue appears homogeneous [Figs. 1(a) and (b)]. Several vertical stripes of high LIV (green) signal around the renal cortex region are observed in the cross-sectional LIV [arrowheads, Fig. 1(c)]. The LIV signals in the vertical tail might be an artifact caused by the high temporal fluctuation of the OCT signal at the top of the structure. Similar vertical artifact appearance was also found in our previous *exvivo* liver study^30^. In the LIV *en face* slice, small structural formations (yellow arrows) and some spatially isolated high-LIV spots (white arrowheads) are visible [Fig. 1(d)].

The cross-sectional Fast-DOCT shows several white hyper-Fast-DOCT (high decorrelation) spots, and no tail artifacts are observable [Fig. 1(e)]. Note that the regions of the hyper-Fast-DOCT spots correlated well with the LIV image [Figs. 1(c) and (e)]. In the magnified view of the Fast-DOCT cross-section (rectangular box), we can identify two closely spaced white spots [Fig. 1(j)] that cannot be recognized in the LIV because of the tail artifact disturbance [Fig. 1(h)]. The *en face* slice of the Fast-DOCT reveals several small white spots that can be identified more clearly in the magnified view [Fig. 1(k)]. Note that the hyper-Fast-DOCT spots are smaller and fewer than the high-LIV spots.

After the dynamic OCT measurement, we imaged the same renal tissue stained with hematoxylin and eosin (H&E) for histology. In the histological micrographs, we can observe several small circular structures surrounded by white space [exemplified by black arrow, Fig. 1(g)]. These structures are the glomeruli of the renal tissue surrounded by Bowman’s capsules^34^. In addition to the glomeruli, renal tissue is comprised of convoluted renal tubule structures, which can be identified in the histology [yellow arrows, Fig. 1(g)].

The reconstructed slab average projection images from the three-dimensional data of the dB-scale OCT intensity, LIV, and Fast-DOCT are displayed in Fig. 2. The slab projection images are from the tissue surface to a depth of 724 μm (see the Methods section for details). The regions of high-LIV signals in Figs. 1(c) and (d) form pipe-like structures in the LIV projection [Fig. 2(c)]. Here the field of view (FOV) is 6 mm × 6 mm. A similar appearance was found in the smaller-FOV measurement (3 mm × 3 mm) [Fig. 2(d)]. No such pipe-like structures were observed in the Fast-DOCT projections for either of the FOVs [Fig. 2(e, f)]. The OCT projection also does not reveal these pipe-like structures, and shows a granular pattern [Fig. 2(a, b)]. The black region at the bottom of the projection images is an artifact [yellow arrow, Fig. 2(a)]. No tissue is located within the measured region, and hence the segmentation did not work and this region appears black.

**Figure 2.**
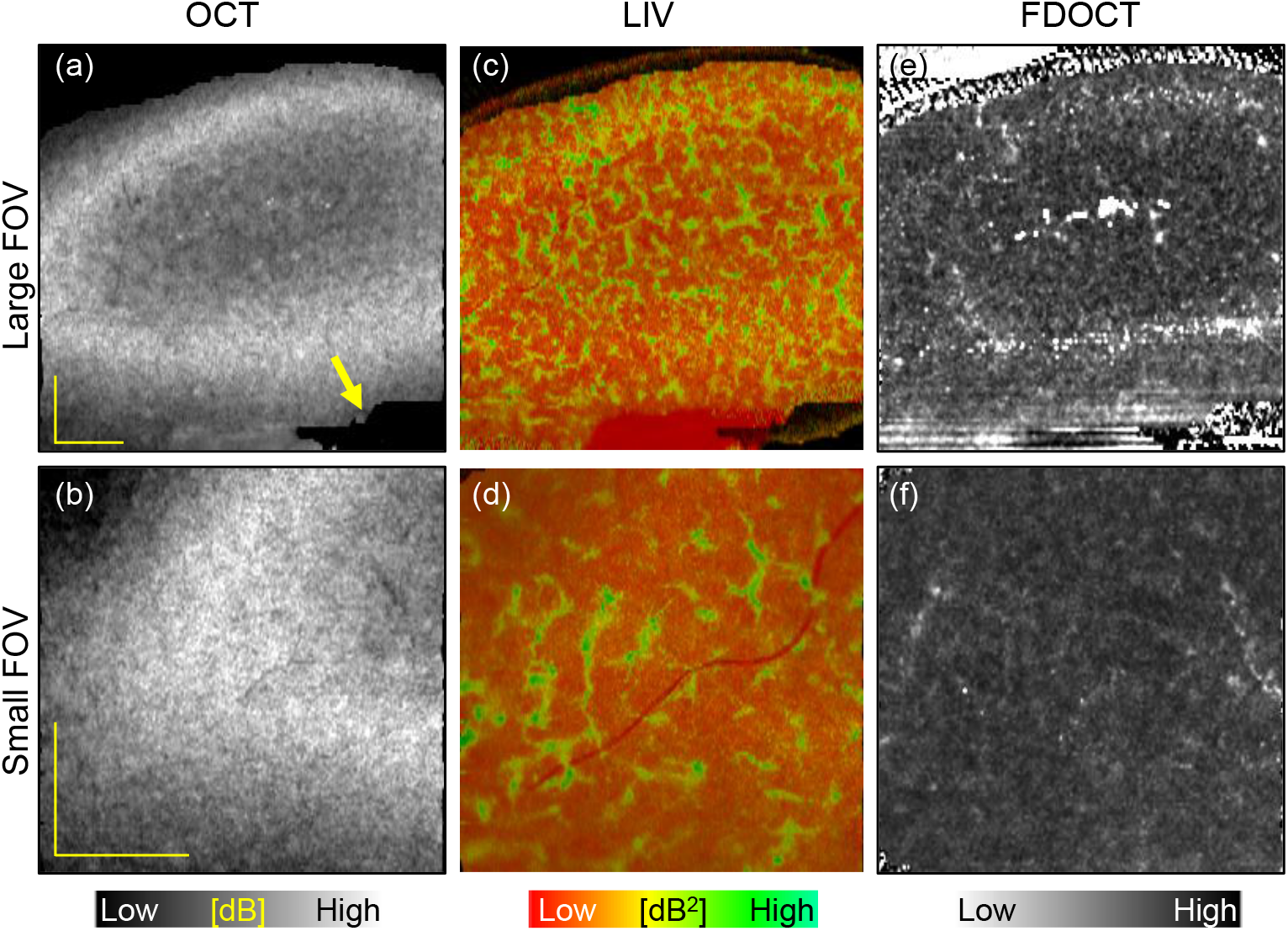
Slab average projection imaging of a normal mouse kidney. (a, b) OCT projections, (c, d) LIV projections, and (e, f) Fast-DOCT projections for large (6 mm × 6 mm, top row) and small (3 mm × 3 mm, bottom row) fields of view. LIV: log-intensity variance; Fast-DOCT: fast dynamic OCT. All scale bars indicate 1 mm.

To confirm whether the dynamics imaging method can detect this structural appearance in other normal renal tissues, we imaged three additional normal kidney samples. Representative LIV and Fast-DOCT imaging of another normal renal tissue sample is illustrated in Fig. 3. The slab projection images of the LIV and Fast-DOCT are displayed in the same arrangement as Fig. 2. Similar to the previous sample, the LIV projection reveals high-contrast pipe-like structures in both FOVs. In this case, both the OCT and Fast-DOCT projection images also reveal these structures. In the OCT projection these structures appear as hypo-scattering regions, whereas in the Fast-DOCT projection they appear as hyper-Fast-DOCT convoluted structures. In the narrower FOV measurement (3 mm × 3 mm), these structures appear more clearly in the OCT and Fast-DOCT projections. It is noteworthy that although the LIV projection contains more detailed functional structures, a similar appearance can also be found in the Fast-DOCT image. We also note that the Fast-DOCT structures correlate well with the regions of the LIV structures, whereas the hypo-scattering structures of the OCT correlate moderately with the LIV and Fast-DOCT. As we discuss later in the Discussion section, these pipe-like structures are the renal tubules of the kidney tissue. In addition, we also explain why these structures appear in the Fast-DOCT image for this case, but not in the previous case.

**Figure 3.**
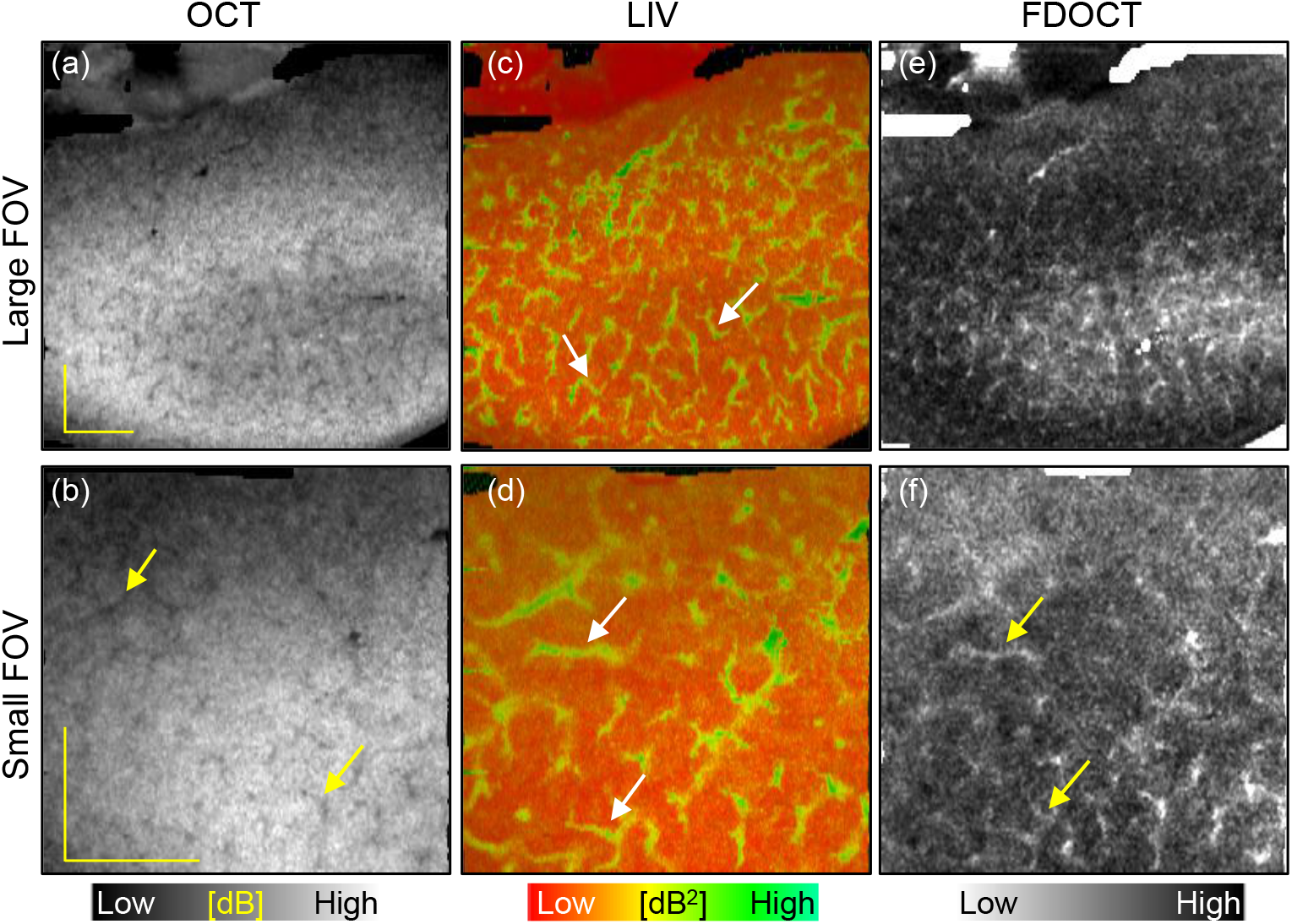
Slab average projection imaging of another normal mouse kidney sample. (a, b) OCT projections, (c, d) LIV projections, and (e, f) Fast-DOCT projections for large (6 mm × 6 mm, first row) and small (3 mm × 3 mm, second row) fields of view. The arrows indicate the pipe-like structures in the renal tissue. LIV: log-intensity variance; Fast-DOCT: fast dynamic OCT. All scale bars indicate 1 mm.

### Label-free dynamics imaging of obstructed kidney models

We take the advantage of label-free functional imaging methods to explore whether we can detect any difference in the morphological renal structural changes between normal kidney and kidney subject to obstruction for several days. We imaged 1-week and 2-week obstructed kidney models using the dynamic OCT microscope and then reconstructed the LIV and Fast-DOCT images from the time-sequential OCT frames. Details of the obstructed kidney model is described later in the Methods section.

Figures 4 and 5 display the OCT, LIV, Fast-DOCT, and histology images of 1-week and 2-week obstructed kidney models,respectively. In both obstructed kidneys, the OCT intensity appeared homogeneous, similar to the normal kidney [Fig. 4(a, b) and Fig. 5(a, b)]. A high-LIV superficial layer (green appearance) at the border of the renal surface region was observed in the cross-sectional and *en face* LIV images of both 1-week and 2-week obstructed kidneys [white arrowheads, Fig. 4(c, d) and Fig. 5(c, d)]. Note that, such a superficial layer like this did not appear in the normal renal tissue. Furthermore, the vertical tails with high LIV signals were also not found in either of the obstructed kidney models. In the 1-week obstructed kidney, two small green isolated dots can be identified in the LIV cross-section [see the rectangular box in Fig. 4(c) and the magnified image Fig. 4(h)]. The LIV *en face* slice of the 1-week model reveals a few small structures similar to those of normal kidneys [black arrows, Fig. 4(d)]. In contrast, these high LIV structures were not observed in the LIV image of the 2-week obstructed kidney model [Fig. 5(c, d)]. Note that the high LIV signal at the center of the *en face* slice of the 2-week obstructed model [white-dot circle, Fig. 5(d)] is an artifact caused by the strong reflection from the tissue surface.

**Figure 4.**
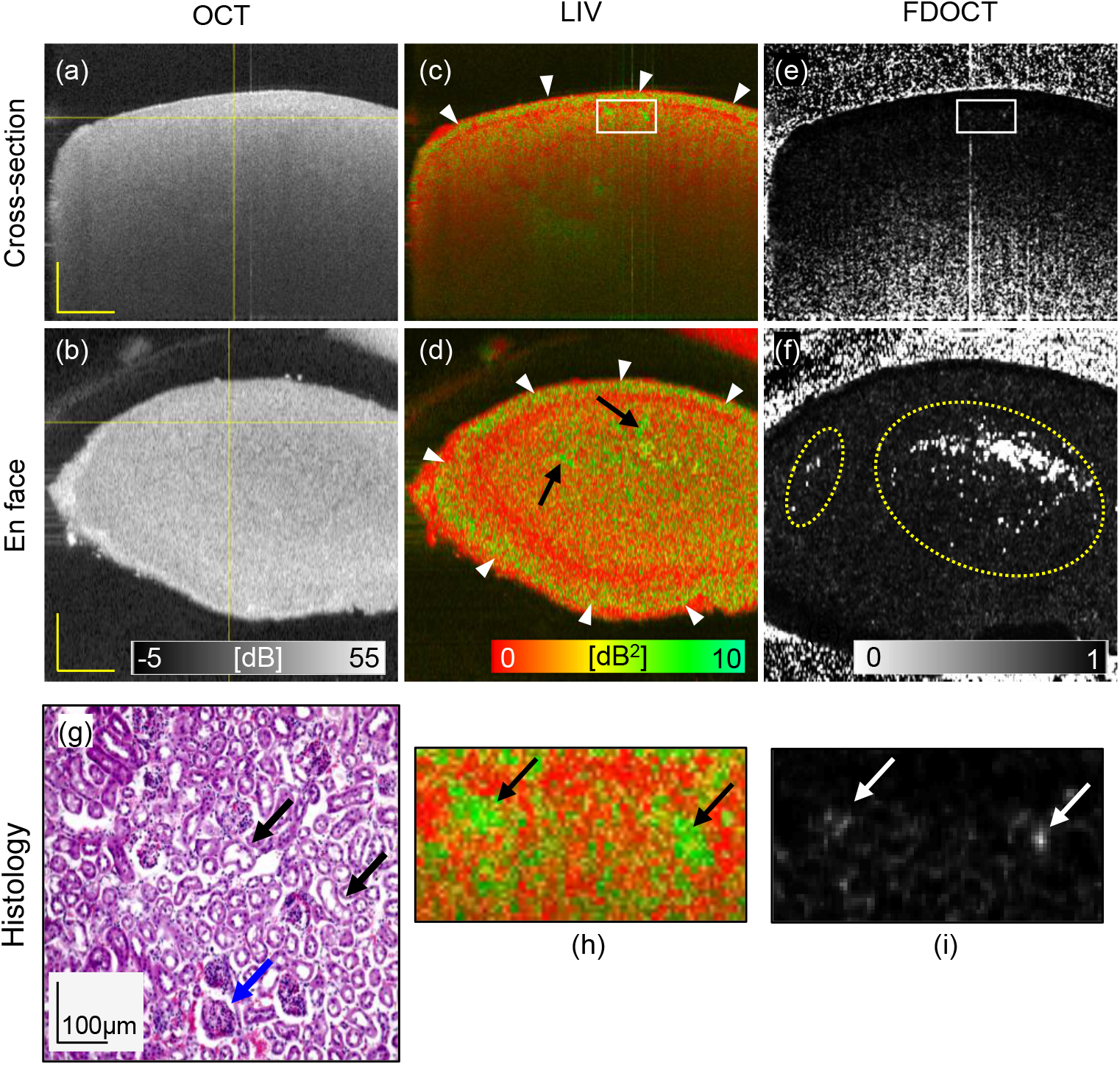
Dynamic OCT imaging of a 1-week obstructed kidney model for 6 mm × 6 mm field of view. Cross-sections from (a)scattering OCT, (c) LIV, and (e) Fast-DOCT imaging; (b, d, f) *En face* slices of the OCT, LIV, and Fast-DOCT images at the depth location indicated by the horizontal line in (a); (g) H&E stained histological micrograph; (h, i) Magnified images of the LIV and Fast-DOCT cross-sections at the region indicated by the rectangular boxes in (c) and (e). The blue arrow in (g) indicates a glomerulus and the black arrows indicate renal tubules of the kidney tissue. LIV: log-intensity variance; Fast-DOCT: fast dynamic OCT. Scale bar: 1 mm.

**Figure 5.**
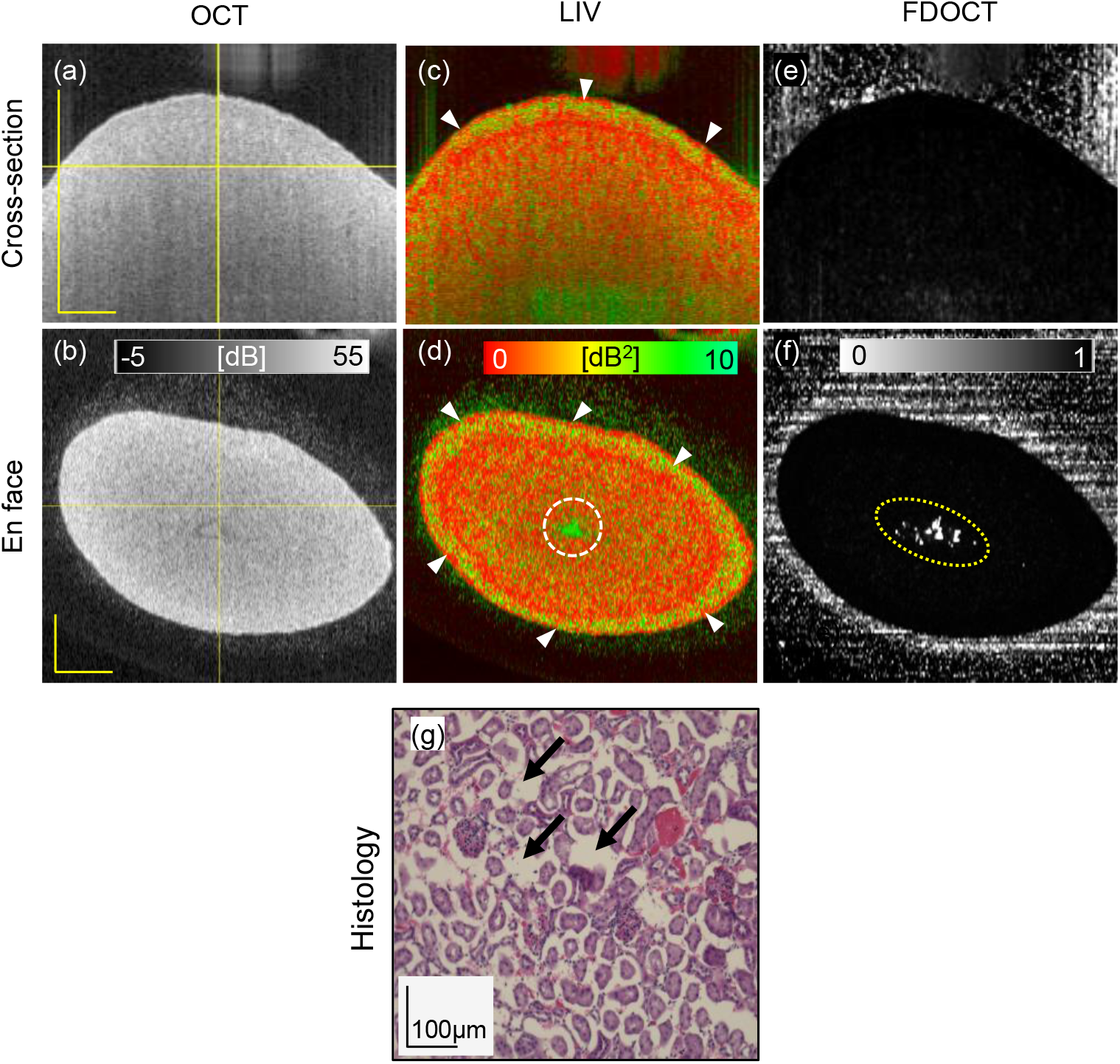
Dynamic OCT imaging of a 2-week obstructed kidney model for 6 mm × 6 mm field of view. Cross-sections from(a) scattering OCT, (c) LIV, and (e) Fast-DOCT imaging; (b, d, f) *En face* slices of the OCT, LIV, and Fast-DOCT images at the depth location indicated by the horizontal line in (a); (g) H&E stained histological micrograph; The circles in (d, f) indicate artifact due to the specular reflection from the surface. LIV: log-intensity variance; Fast-DOCT: fast dynamic OCT. Scale bar: 1 mm.

The cross-sectional Fast-DOCT of the 1-week obstructed kidney also reveals hyper-Fast-DOCT spots that correlate well with the position of the high LIV [Fig. 4(e) and magnified image Fig. 4(i)]. These white Fast-DOCT spots appear as relatively low contrast in the 1-week model in comparison with the normal kidney. On the other hand, the white dots in the *en face* Fast-DOCT images for both 1-week and 2-week obstructed kidneys [yellow-dot circle, Fig. 4(f) and Fig. 5(f)] are artifacts, and are due to the strong reflection from the tissue surface. In the 2-week obstructed kidney model, the Fast-DOCT image [Fig. 5(e) and (f)] appeared as completely black, except for the artifacts.

The glomeruli and renal tubules can be identified in the histology images of both 1-week and 2-week obstructed kidney models [Fig. 4(g) and Fig. 5(g)]. Histological micrographs of the 2-week model show some structural damage and dilation in the renal tissue [Black arrows, Fig. 5(g)].

### Comparison of dynamics imaging between contralateral non-obstructed and obstructed kidneys

We further explored the possibility of imaging the contralateral non-obstructed and obstructed kidneys of a single mouse. The left kidney was obstructed by tying its ureter with a nylon thread, and is referred to as the obstructed kidney, while the right kidney on the opposite side is referred to as the non-obstructed kidney.

A comparison between the slab average projection images of the contralateral non-obstructed and obstructed kidneys is displayed in Fig. 6. The OCT projection of the obstructed kidney shows a granular hypo-scattering pattern [Fig. 6(b)] that is not identical to the non-obstructed kidney [Fig. 6(a)]. The LIV projection of the non-obstructed kidney shows similar pipe-like structures to those found in normal kidney [Fig. 6(c)]. Again, these structures are not visible in the obstructed kidney, similar to the 2-week obstructed model [Fig. 6(d)]. The Fast-DOCT projection images do not show these structures in either kidney [Fig. 6(e) and (f)]. The absence of the pipe-like structures from the LIV images of the obstructed kidney may indicate the loss of functionality of the renal tubules due to the ureter obstruction, as discussed later in the Discussion section.

**Figure 6.**
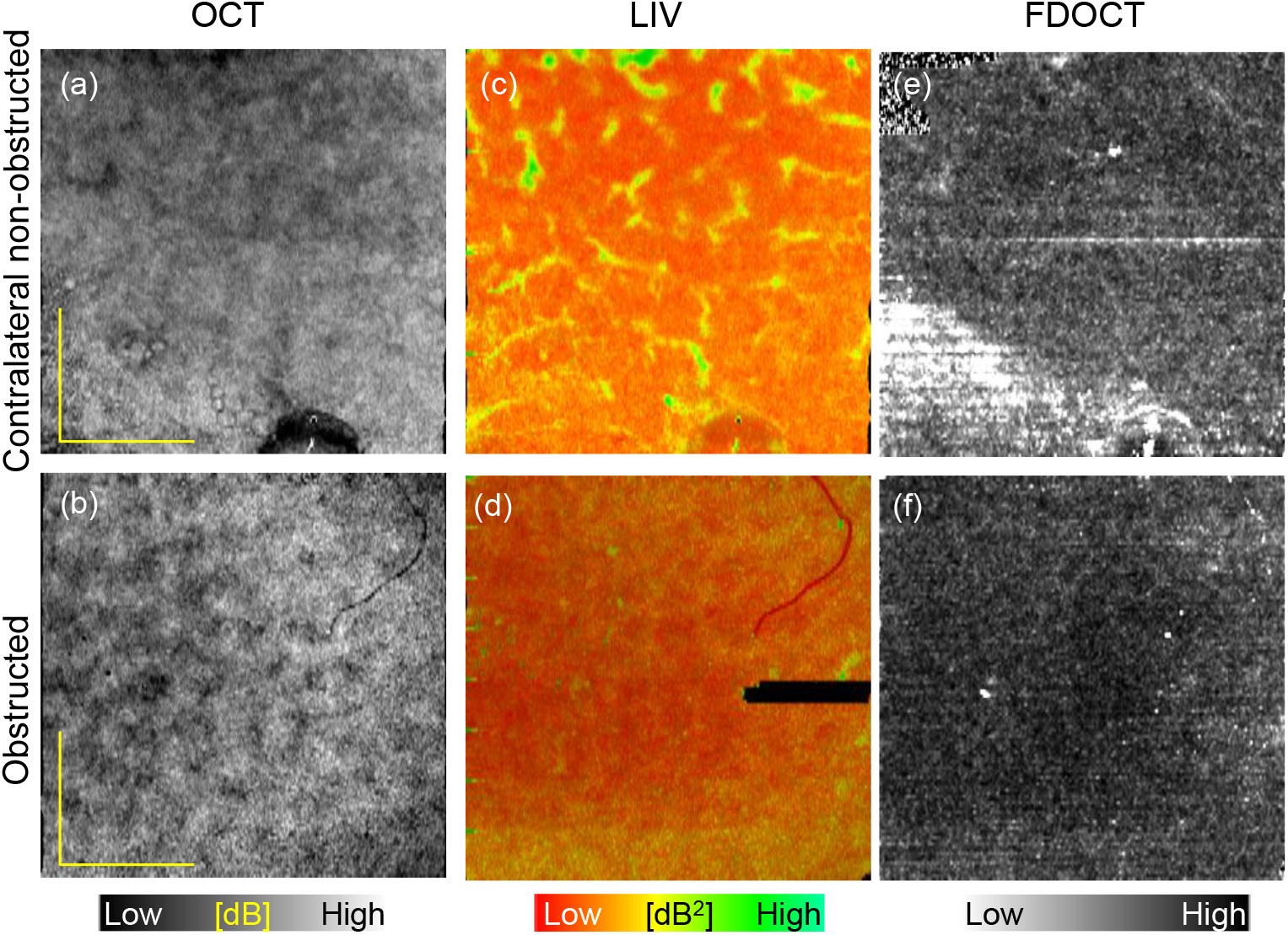
Slab average projection imaging of contralateral non-obstructed (first row) and obstructed kidney (second row) models. (a, b) OCT projections, (c, d) LIV projections, and (e, f) Fast-DOCT projections for 3 mm × 3 mm field of view measurement. LIV: log-intensity variance; Fast-DOCT: fast dynamic OCT. All scale bars indicate 1 mm.

## Discussion

Here we first show the pipe-like structures in the DOCT (LIV and Fast-DOCT) projection image of the normal kidney [Figs. 1 - 3] are renal tubules. The nephron is the microscopic structural and functional unit of the kidney and consists of two parts: renal corpuscles and renal tubules [Fig. 7(a)]^1^. The renal corpuscles contain cluster of capillaries, called the glomeruli, which are surrounded by Bowman’s capsules. The renal tubules consist of the proximal convoluted tubule (PCT), the distal convoluted tubule (DCT), and the loop of Henle. Several previous studies demonstrated renal tubular imaging in animal kidneys using OCT. For example, Chen *et al*. demonstrated fine superficial renal tubular structural imaging in Munich-Wistar-strain rat kidney *ex vivo* using a high-resolution OCT (axial - 3.3 μm, and lateral - 6 μm) with a broadband laser light source with a center wavelength of 1.3 μm)^14^. Three-dimensional morphology of the renal tubule was also illustrated in Munich-Wistar-strain kidney *in vivo*^15^ by the same group. In both studies, the renal tubule appeared as a lumen in the OCT cross-section and as a hypo-scattering structure in the *en face* image, similar to ours [Fig. 3(a, b)]. Thus, we can conclude that the low-backscattering structures in the OCT are renal tubules.

**Figure 7.**
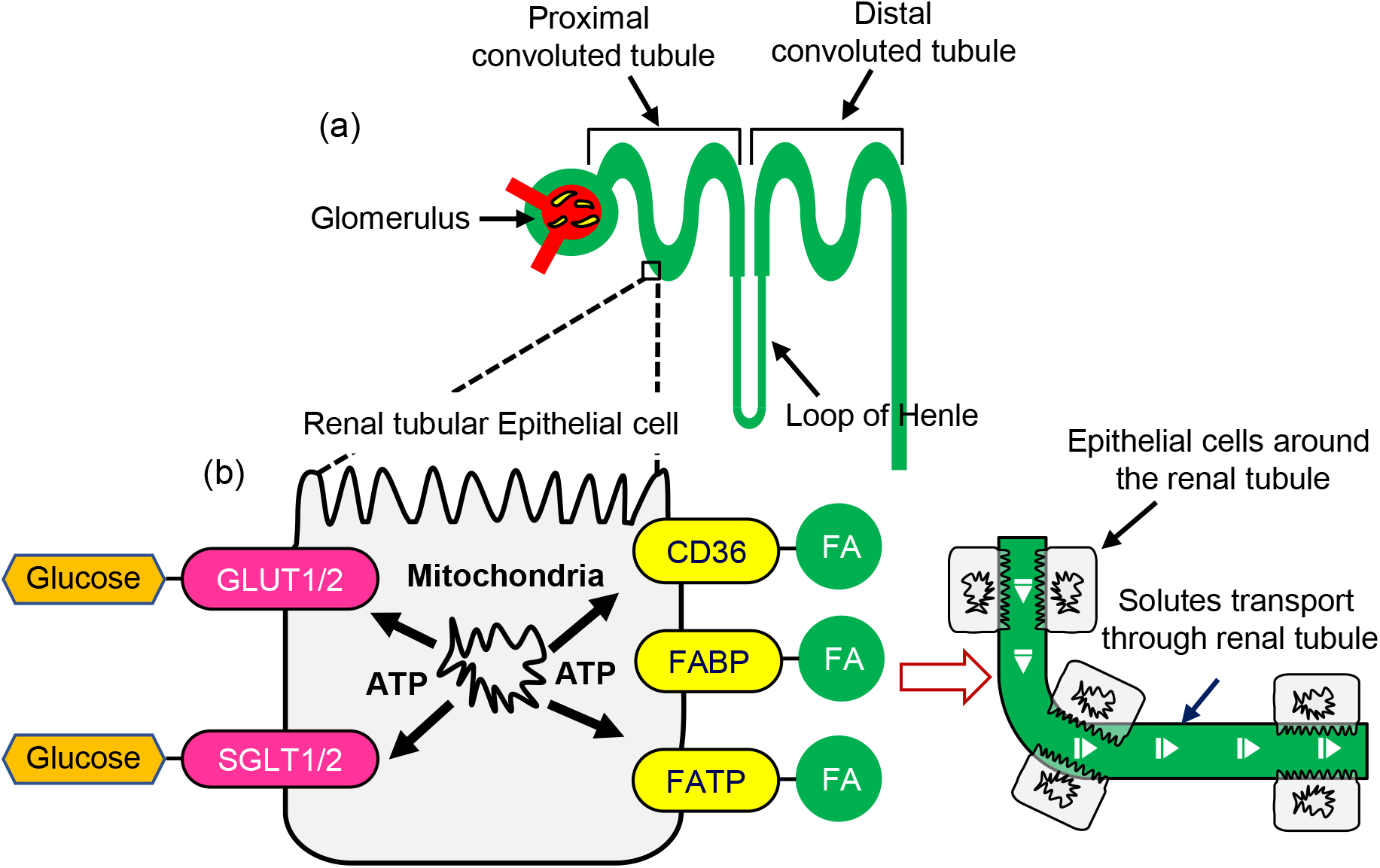
(a) Anatomical structure of the nephron and (b) schematic of the function within the renal tubule. GLUT1/2 and SGLT1/2: glucose transporters; CD36: receptor; FABP: fatty acid-binding proteins; FATP: fatty acid transport proteins; FA: fatty acid.

We determined the diameter of these pipe-like structures from the OCT projection image and compared it with previous OCT measurements^14,15^. We found that the diameter of this structure was around 20 – 25 μm, which is close to that described in previous studies (30 – 40 μm). In addition, the structures observed in the DOCT projection images have a similar convoluted appearance to those observed in previous OCT studies, and also in the corresponding histology. Therefore, we believe that the pipe-like structures in the DOCT projection images are renal tubules. Note that the diameter found from the LIV image is slightly larger than that from the OCT, around 38 – 51 μm. We will discuss the reason for this difference in a later paragraph of this Discussion section.

There might be some curiosity regarding why the pipe-like structures are not clearly visible in the standard OCT intensity in this study, while they were clearly visible in previous studies^14,15^. This can be explained by two reasons. First, the resolution of the OCT system used in this study (18.1 μm laterally and 14 μm axially) is lower than that of previous OCT studies of the kidney, making it difficult to visualize renal tubule structures in the OCT intensity image. Second, the current study used mouse kidney tissue, whereas previous studies used Munich-Wistar strain rat, which is known to have more-visible fine superficial renal tubules. In the future, it might be possible to visualize such renal tubular structures in the LIV as well as in the OCT with Contralateral non-obstructe our high-resolution OCT^35^ or by imaging the Munich-Wistar strain rat.

We suspect that the high DOCT signal in the pipe-like structures originates from active renal tubular epithelial cells (RTECs) around the renal tubule. A schematic of renal tubule function is shown in Fig. 7. The kidney relies on the renal tubules to reabsorb most of the filtered water and solutes, including glucose and fatty acids. The renal tubule is made up of epithelial cells that are responsible for the active transport process through the renal tubule. RTECs have mitochondria that provide an excessive amount of ATP energy to glucose and fatty acid transporters, and hence RTECs are functionally active. Glucose is transported through the SGLT1/2 and GLUT1/2 transporters^36^. Fatty acid transport through the renal tubules by the CD36 receptors, fatty acid-binding proteins (FABP), and fatty acid transport proteins (FATP)^9,37^. We believe that after we sacrifice the mouse, the active transport process stopped in the renal tubule, but the RTECs around the renal tubules were still highly active. The active RTECs can cause intercellular motion resulting in speckle fluctuation, and hence could result in a high DOCT signal around the renal tubule.

We mentioned earlier that the diameter of the pipe-like convoluted structures was larger in LIV than in the OCT image. This can be explained by the previous discussion that what we are observing in the LIV image might be the functional activity of the RTECs around the renal tubule. As the high LIV signals are from the surrounding tissue around the renal tubule, it seems plausible that the diameter of these structures is higher in LIV compared to OCT. Note that, in our previous study on mouse liver^29^, we found similar surrounding tissue activity. Specifically, we observed high activity just beneath the liver vessel using LIV imaging, which was due to the high metabolism of the periportal and pericentral regions of the liver microvasculature.

Next, we discuss our findings in the obstructed kidney models. The anatomical pipe-like structures that were observed in the LIV of normal kidneys were almost not found in the 1-week model and were completely absent from the 2-week model. Literature suggests that obstruction of the ureter for a certain period results in renal hemodynamic and metabolic changes, followed by tubular injury, and finally apoptosis or necrosis^38^. During the apoptosis or necrosis process, the function of the renal tubule can gradually alter and is finally lost due to cell death. The histology of the 2-week obstructed model also suggests renal structural damage and dilation, which further highlight the process of renal injury due to obstruction. The absence of the pipe-like structures in the LIV and Fast-DOCT images in the obstructed models may therefore indicate the loss of renal tubule function due to the effect of obstruction.

Instead of any anatomical structures, we found a high-LIV superficial layer around the peri-renal cortex region in both obstructed kidney models [Figs. 4 and 5]. Similar results to those in Figs. 4 and 5 were obtained for all obstructed kidne through renal tubul model samples. The histology images of these samples did not reveal any changes around the peri-renal cortex region and appeared similar to the those of normal kidney. To the best of our knowledge, such an appearance was not reported previously by fluorescence microscopy imaging.

Literature suggests that the obstruction of the ureter for a certain period induces renal inflammation^39,40^. Bai *et al*. reported that several pro-inflammatory cytokine IL-1*β* and transforming growth factor-*β*1 (TGF-*β*1) are recruited in the renal environment during the obstruction^41^. Due to regulatory mechanisms involved during renal inflammation, these inflammatory cells (IL-1*β* and TGF-*β*1) can be highly metabolically active. We suspect that during obstruction, renal inflammation occurs near the peri-renal cortex region, and since the inflammatory cells are active, they can produce speckle fluctuation and result in a high LIV signal. Although we discuss a possible cause, it is not conclusive, and such a peri-renal layer appearance on the LIV image is worthy of further investigation.

In this study, we employed two different DOCT algorithms to access the slow and fast functional activity of renal tubules; one based on logarithmic intensity variance and one on complex correlation analysis of time-sequence OCT intensity. In our pilot study, we detected several vertical artifacts in the LIV cross-section of normal kidneys, similar to the OCT angiography (OCTA) projection artifacts of retinal imaging^42^. This artifact suggests that there might be some structure in the renal tissue that may exhibit relatively fast motion. The LIV contrast is too sensitive to visualize this fast motion in the tissue. On the other hand, our previous study also found that when the time window is small (milliseconds), LIV cannot visualize the fine high dynamics structures^18^. To overcome the above limitation of LIV, we employed complex-correlation-based Fast-DOCT contrast in addition to LIV to visualize structures with high dynamics, and imaging with this contrast did not exhibit any tailed artifacts.

We also note that LIV and Fast-DOCT were reconstructed from two volumes using two different scanning protocols (details are in the Methods section). In the future, a new scanning protocol may overcome this issue allowing both contrasts to be reconstructed from a single volume^43^.

As we showed previously, that the Fast-DOCT projection reveals pipe-like structures in some renal tissues [Fig. 3(f)] but not in others [Fig. 2(f)]. This discrepancy may be attributed to differences in the measurement time between samples. Namely, several mice were sacrificed at a time, and then the extracted kidneys were sequentially measured one by one using the OCT system and raster scanning protocols (see the details in the Methods section). Due to the *ex vivo* study, the function of the kidneys could have gradually decreased even if they were kept in the cultured medium and this could possibly be the reason that not all the kidneys revealed such pipe-like structures. Although the Fast-DOCT projections did not reveal the pipe-like structure in some normal renal tissues, we could still observe several isolated white spots in both cross-sections and *en face* slices. In contrast, the same white hyper-Fast-DOCT spots were not found completely in any cross-section or *en face* image of the obstructed kidney models.

## Conclusion

The functional activity of normal kidney, a 1-week obstructed kidney model, and a 2-week obstructed model, were investigated three-dimensionally and without any labeling by volumetric dynamic OCT technique. LIV and Fast-DOCT imaging successfully visualized superficial pipe-like functional structures in the normal kidney that may correspond to renal tubules. These structures were hardly visible in the 1-week obstructed model and were completely absent in the 2-week obstructed model. The absence of pipe-like structures in obstructed kidney models may suggest the loss of renal tubular function due to the effect of obstruction. Instead of any anatomical structures, a circular shell at the peri-renal cortex region was visualized in the LIV in both 1-week and 2-week obstructed models that was not found in normal kidney. In conclusion, dynamic OCT imaging can be used to visualize the functional activity of renal tubules without any exogenous labeling, and hence can be used as a tool to assess and investigate kidney injury in animal models.

## Methods

### Obstructed mouse kidney model and study design

Experiments were performed on normal mouse kidneys and obstructed mouse kidney models *ex vivo*. Eleven C57BL/6 mice at an age of 4 – 5 months were used in the experiments. The mice were housed in normal conditions under a 12-hour light/dark cycle.

For the normal mouse kidney study, the mice were sacrificed via cervical dislocation and the kidneys were extracted and placed in a petri dish. After dissection, the kidney was submerged in Dulbecco’s modified eagle medium (DMEM, Sigma-Aldrich, MA) supplemented with 10% fetal bovine serum (Sigma-Aldrich, MA) and penicillin/streptomycin (Gibco, MA), and kept at room temperature during the experiment. OCT imaging was performed almost 1-hour after the dissection. After the OCT measurement, the same renal tissue was fixed in paraformaldehyde and stained with H&E.

For the obstructed mouse kidney model, mice were anesthetized by a combination of anesthetics [medetomidine (Nippon Zenyaku Kogyo), midazolam (Astellas Pharmaceuticals), and butorphanol (Meiji Seka Pharma)] and the left ureter was then exposed by a dorsal incision. The left ureter was obstructed by ligation with nylon sutures and the right ureter was left as it was. Following the surgical procedure, mice were injected with anesthetic antagonist, Atipamezole (Nippon Zenyaku Kogyo). The ligation in the left ureter was applied for either 7 days or 14 days. The kidney of the ligated left ureter is referred to as the obstructed kidney and the contralateral right kidney as the non-obstructed kidney. Obstructed kidney subjected to ureter ligation for 7 days is referred to as the 1-week obstructed kidney model, while kidney subjected to 14 days of ligation is referred to as the 2-week obstructed kidney model. After 7 or 14 days, the mice were sacrificed and the kidneys were extracted and placed in a petri dish. Similar to the previous study on normal kidney, the obstructed and non-obstructed kidneys were submerged in DMEM cultured medium and kept at room temperature during the OCT measurement. H&E staining histology was performed after the OCT measurement.

All animal experiments were performed in accordance with the animal study guidelines of the University of Tsukuba. All experimental protocols involving mice were approved by the Institutional Animal Care and Use Committee (IACUC) of the University of Tsukuba. The present study was designed, performed, and reported according to the principles of ARRIVE (Animal Research: Reporting of In vivo Experiments) guidelines.

### Swept-source OCT system

A custom-made swept-source-based OCT device was used in this study^32,33^. The system uses a MEMS-based wavelength sweeping laser (AXP50124-8, Axun Technologies, MA) with a scanning frequency of 50 kHz, a center wavelength of 1.3 μm, and a depth resolution of 14 μm in tissue. The system was used in combination with a scan lens (LSM03, Thorlabs Inc., NJ) having an effective focal length of 36 mm, and the effective numerical aperture (NA) of the probe optics is 0.048. The lateral resolution is 18.1 μm and the lateral pixel separation is 1.95 μm. The probe power on the sample was 12 mW and the system sensitivity is 104 dB. More detailed descriptions of the system specifications are available elsewhere^32,33^.

### Dynamic OCT imaging algorithms

Two different signal processing algorithms were used to visualize slow (second-scale time window) and fast (millisecond-scale time window) dynamics. One is LIV which is sensitive to slow dynamics, and another is complex-correlation-based Fast-DOCT which is sensitive to fast millisecond-scale tissue dynamics.

LIV is the contrast for measuring the OCT signal fluctuation magnitude. The LIV signal was computed using the variance of the time-sequence logarithmic (dB)-scale OCT intensity signal^18^.

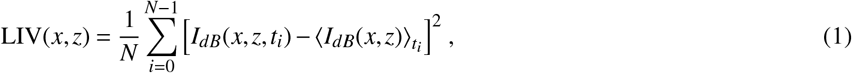

where *I*_*dB*_ is the dB-scale OCT intensity, *x* and *z* are the lateral and axial positions, respectively, *t*_*i*_ is the acquisition time point of the *i*-th frame, 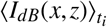 is the averaged dB-scale OCT intensity over *t*_*i*_, and *N* is the number of frames.

The LIV images are shown by pseudo-color composite image as described in Section 2.3 of Ref. 18. In this composite image, the Hue (H channel) is the LIV, the pixel brightness (V channel) is the dB-scale OCT intensity, and the saturation (S channel) is set to 1 for all pixels. In the LIV image, regions with high dynamics appear as green and regions with low dynamics appear as red.

Fast-DOCT is based on OCT angiography (OCTA) technology, and is computed through complex correlation analysis among the four repeated frames with noise correction^31^; it is identical to the complex correlation-based OCTA. Specifically, we used noise-corrected OCTA, which is a complex decorrelation with a noise offset correction.

### Scanning protocols

Two different scanning protocols were used to obtain volumetric LIV and Fast-DOCT images. Three-dimensional LIV was obtained using a repeating raster scan protocol^25^. For the repeating raster scan, the whole volumetric region (covering a 3 mm × 3 mm or 6 mm × 6 mm lateral region) was divided into four blocks, with each block covering 3 mm × 0.75 mm ×or 6 mm ×mm.1.5mm In each block, 32 B-scan locations were first scanned by a raster scan, with each cross-sectional image (i.e., frames) consisting of 512 A-lines. The raster scan was repeated 16 times with an inter-scan time interval of 409.6 ms. Therefore, at each B-scan location, 16 sequential frames were obtained in a 6.12-s time window. All blocks were then sequentially measured to obtain the LIV volume. The final LIV volume consisted of 2048 frames in total, for 128 B-scan locations in the sample. The total acquisition time to capture this volume was 26.2 s.

Fast-DOCT was obtained using another fast raster scanning protocol similar to that of OCTA. Using a fast raster scan, four frames with an interval of 12.8 ms were sequentially acquired at each B-scan location. Therefore, at each B-scan location, four frames were acquired in a 39-ms time window and each frame consisted of 512 A-lines. The final volume was obtained by sequentially scanning 128 B-scan locations in the sample. The volume acquisition time for 512 frames was 6.55 s.

### Axial slab average projection images

To create axial slab average projection images, the tissue surface was first identified from the dB-scale OCT intensity image by applying a surface segmentation algorithm, as described in^33^. Here, the OCT intensity is the average of 16 frames. The depth region from the surface to 100 pixels (724 μm) was extracted. The process was performed for all the frames in a 3D volumetric dataset to form a slab. Next, the slab projection images of OCT, LIV, and Fast-DOCT were obtained by averaging the respective intensity, LIV, and Fast-DOCT in depth (axially), based on the volumetric datasets of OCT, LIV, and Fast-DOCT. Note that the average projection images provide better contrast than the maximum intensity projections.

### Histology

The samples were preserved in 4% paraformaldehyde (PFA) (Nacalai Tesque, Kyoto, Japan) after the OCT measurement. The kidneys were then embedded in paraffin, and thin sections of 12 μm thickness were prepared using a cryosection procedure. Hematoxylin and eosin (H&E) was used to stain the sections, and the H&E-stained sections were imaged under a microscope (BZ-X710, Keyence Corp., Osaka, Japan) with a 10× objective (NA = 0.3).

## Acknowledgments

Funding was provided by Core Research for Evolutional Science and Technology (JPMJCR2105); the Japan Science and Technology Agency (JPMJMI18G8); and the Japan Society for the Promotion of Science (18H01893, 21H01836, 22K04962, 22F22355). Fruitful technical discussions with Arata Miyazawa (Skytechnology) and Antonia Lichtenegger (Medical University of Vienna, Austria) are gratefully acknowledged.

## Author contributions

P.M., S.F., and Y.Y. designed the study. S.F., D.L., and T.H.T prepared the samples and performed surgery to create the obstructed kidney model. P.M. organized the experiments and collected all data. P.M., S.F., K.O., I.A.S., Y.L., S.M., and Y.Y. analyzed and interpreted the data. P.M. wrote the first draft of the paper, and all authors revised the work and approved the final version of the manuscript. S.F. and Y.Y. supervised the work.

